# A comparison of sharpening and dampening accounts of the role of expectation in shaping the neural fidelity of early visual representations

**DOI:** 10.64898/2026.07.05.736624

**Authors:** Reuben Rideaux, Ziyue Hu, Kali Chidley, Martijn A Cloos, D Samuel Schwarzkopf, Jason B Mattingley

**Affiliations:** School of Psychology, The University of Sydney, Camperdown, Australia; Queensland Brain Institute, The University of Queensland, St Lucia, Australia; School of Psychology, The University of Queensland, St Lucia, Australia; Australian Institute for Bioengineering and Nanotechnology, The University of Queensland, St Lucia, Australia; Donders Centre for Cognitive Neuroimaging, Radboud University, Nijmegen, Netherlands; School of Optometry & Vision Science, Waipapa Taumata Rau University of Auckland, New Zealand; Experimental Psychology, University College London, United Kingdom

## Abstract

The natural environment is spatiotemporally structured, and the brain exploits this regularity to predict and prepare for upcoming sensory stimuli. Such predictive processing is thought to increase neural efficiency by reducing metabolic expenditure and altering the fidelity with which newly encountered stimuli are encoded. Competing theoretical frameworks propose this is achieved either through sharpening, whereby expected events are encoded more precisely, or dampening, whereby expected events are suppressed and encoded less precisely. Despite clear, opposing predictions, evidence in humans for each account remains mixed due to methodological and analytical inconsistencies. Here we addressed these issues using probabilistic visual paradigm combined with functional magnetic resonance imaging (fMRI) and electroencephalography (EEG). We used population receptive field (pRF) mapping of fMRI data and inverted encoding of EEG data to compare the fidelity and timecourse of activity in visual areas in response to expected, unexpected, and random stimuli. Both methods produced a consistent pattern of results. Post hoc analysis of EEG data revealed that the apparent effect of expectancy was better explained by local spatiotemporal stimulus properties than the global expectancy manipulation. Although this pattern resembled sensory adaptation, it was more consistent with an expectation of temporal stability combined with dampening, in which both the aggregate response to expected features and their representational fidelity are suppressed. Taken together, our findings suggest that predictive processing may operate through dampening, with ecological advantages for high-fidelity encoding of unexpected sensory events.

## INTRODUCTION

The brain is sensitive to the spatial and temporal structure inherent in the natural environment, and can use this structure to predict and prepare for upcoming sensory inputs. For example, moving objects typically follow predictable trajectories that can be anticipated to support the effectiveness of motor behaviours ranging from smooth pursuit eye movements and saccades to collision avoidance and interception (Wolpert & Flanagan, 2001). Anticipation of sensory events may also be used to support neural efficiency, by reducing metabolic expenditure and increasing the fidelity with which sensory events are encoded (Barlow, 1961; Friston, 2005; Rao & Ballard, 1999).

Two popular frameworks - sharpening and dampening – have been proposed to explain the mechanism by which expectation shapes sensory processing. While the two explanations agree that metabolic expenditure is reduced in response to expected sensory events, they differ in the means through which this is achieved, and offer opposing predictions for the fidelity of expected and unexpected stimulus encoding. The sharpening account claims that neurons tuned to near-to-expected features are suppressed, resulting in expected events being encoded more precisely (Kok et al., 2012; Teufel et al., 2018; Yon et al., 2018). By contrast, the dampening account asserts that expected features are suppressed, resulting in expected events being encoded less precisely (Lee & Rideaux, 2025; Rideaux, Dang, et al., 2025; Tang et al., 2018, 2023). A common approach to experimentally testing the effects of expectancy has been to use probabilistic cues and compare behavioural or neural responses to target stimuli that are either likely or unlikely based on the preceding cue. The sharpening account predicts that probabilistically likely (expected) stimuli will be encoded more sharply than unlikely (unexpected) stimuli, and the dampening account predicts the opposite pattern.

Despite the clear and falsifiable predictions made by the sharpening and dampening frameworks, there is still no consensus as to which account better describes the influence of expectation on the fidelity of perceptual representations. Compelling evidence for both accounts has been provided across a range of experimental designs and neuroimaging methods (Kok et al., 2012; Rideaux, Dang, et al., 2025; Tang et al., 2023, 2023; Teufel et al., 2018; Yon et al., 2018). There are several possible explanations for these seemingly discrepant findings. The effects of expectancy can be challenging to separate from other phenomena, such as sensory adaptation and attention, which can confound results and lead to misattribution of altered sensory processing (den Ouden et al., 2023; Feuerriegel, 2024). Methodologically, biophysical differences in the signals recorded by non-invasive neuroimaging methods, such as functional magnetic resonance imaging (fMRI) and electroencephalography (EEG), may play a role in determining which framework is supported, even within the same experimental design. For example, if sharpening and dampening *both* occur, but at distinct temporal delays or cortical locations (Press et al., 2020; Teufel & Fletcher, 2020), these methods, which vary in their spatiotemporal resolution, may produce conflicting results.

While phenomenological and methodological factors are strong candidates for contributing to empirical discrepancies, another factor that may further obscure consensus are analytical differences, particularly in human studies where representational fidelity is inferred from aggregate or indirect metrics of neural activity. For instance, while the sharpening and dampening accounts make opposing predictions about the sharpness with which expected events are encoded, multivariate decoding accuracy (i.e., comparisons between local patterns of neural activity) is typically used as a proxy of this precision, with higher values interpreted as evidence for more precise representations (Kok et al., 2012; Tang et al., 2018; Yon et al., 2018). Indeed, decoding accuracy is assumed to reflect the discriminability of stimuli, such that if a stimulus is represented more sharply it will be more distinguishable from other stimuli and thus lead to more accurate decoding. However, decoding accuracy also reflects the reliability of signals over multiple measurements, which may change independently of encoding precision. For example, a sensory event may elicit a neural response that is similar to other sensory events within a given feature-space (e.g., orientation), but if the reliability of the response across repeated presentations is increased, then decoding accuracy will improve, despite the sharpness of the representation remaining unchanged (Stehr et al., 2023). Thus, using decoding accuracy to test whether expected events are encoded more or less sharply has limitations that may contribute to the current lack of clarity around the influence of expectation on sensory encoding.

An alternative analytic approach, which is commonly used to characterize the spatial selectivity of neuronal populations in visual cortex, and which may better distinguish between response reliability and tuning width, is population receptive field (pRF) mapping (Dumoulin & Wandell, 2008). This approach extends the classical notion of a single neuron’s receptive field—the region of sensory space where stimuli elicit responses—to the voxel level measurable with fMRI. In this framework, the response of a voxel is modelled as arising from the aggregate activity of many neurons with overlapping receptive fields. During pRF mapping, participants view stimuli (e.g., drifting bars, wedges, or expanding/contracting rings) that systematically traverse the visual field while blood-oxygen level-dependent (BOLD) responses are recorded. A computational model is then fit to the voxel time series, typically assuming a parametric receptive field shape (often a 2D Gaussian defined by position and size). By estimating the model parameters that best explain the observed responses, the visual field location and spatial extent over which each voxel responds can be inferred. Thus, in contrast to decoding accuracy, pRF mapping provides a more direct measure of representational sharpness (specifically, pRF width), which can be used to adjudicate between sharpening and dampening accounts of predictive processing.

Here we used pRF mapping to compare the tuning width of receptive fields in visual cortex in response to otherwise identical visual stimuli expected, unexpected, or random. Data from ten participants were collected over three scanning sessions per individual (30 scanning sessions in total) using ultra-high field (7T) fMRI. Data from a further 24 participants were collected using EEG, using the same experimental paradigm, to exploit its superior temporal resolution. EEG data were analyzed using inverted encoding, which reconstructs stimulus features by first modeling how neural responses encode those features and then mathematically inverting that encoding model to estimate the feature representation from population activity patterns. Stimuli comprised checkerboard wedges presented at different polar angles, and their expectancy (expected, unexpected, no expectation) was manipulated through probabilistic cueing (high, low, and equal probability). The aim of the study was to compare responses to stimuli in the three expectancy conditions in both fMRI and EEG datasets to adjudicate between sharpening and dampening accounts of predictive processing.

## METHODS

### Participants

Ten human adults were recruited from The University of Queensland for the fMRI experiment (6 females; average age, 27.1±5.0 years), and 24 were recruited from the University of Sydney for the EEG experiment (18 females; average age, 23.4±6.0 years). Sample sizes were based on previous related fMRI (Infanti & Schwarzkopf, 2020) and EEG experiments (Rideaux, 2024). Participants were neurotypical (by self-report), had normal or corrected-to-normal vision (assessed using a standard Snellen eye chart), and naïve to the aims of the study. The fMRI and EEG experiment protocols were approved by Human Ethics Committees at The University of Queensland (2020/HE003101) and The University of Sydney (2023/HE000072).

### Stimuli and procedure

Stimuli comprised monochrome checkerboard wedges, presented on a mid-grey background, which subtended a visual angle of 12° from the central fixation point, with an arc of 22.5° (**Fig. 1a**). Wedges were presented for 750 ms at one of 16 possible linearly spaced polar angles between 0-360°, and alternated polarity at ∼6.6 Hz. A black dot (radius, 0.1°) was presented centrally throughout the experiment, to support stable fixation. In each trial, a pair of wedges was presented sequentially, with pairs separated by a blank inter-pair-interval of 1500 ms. On 80% of trials, the second wedge in each pair was presented at the polar opposite location to the first (high probability, “expected” stimuli); in the remaining trials, the second wedge was presented at any of the remaining 16 locations, excluding the location of the first wedge and its polar opposite (low probability, “unexpected” stimuli; **Fig. 1b**). To establish expectations in participants, the first eight trials of each block were always high probability stimuli. Trials were presented in blocks of 240 (∼8 min), and each experimental session comprised five blocks. In the fMRI experiment, each participant completed two experimental sessions and one retinotopic mapping session, each ∼1 h in duration. In the EEG experiment, each participant completed a single experimental session (∼1 h). In the fMRI experiment, participants additionally completed an initial familiarization session, which was the same as the experimental sessions, but was undertaken in a behavioural lab without neural recordings. The data from the familiarization session were not included in any of the analyses.

**Figure 1.**
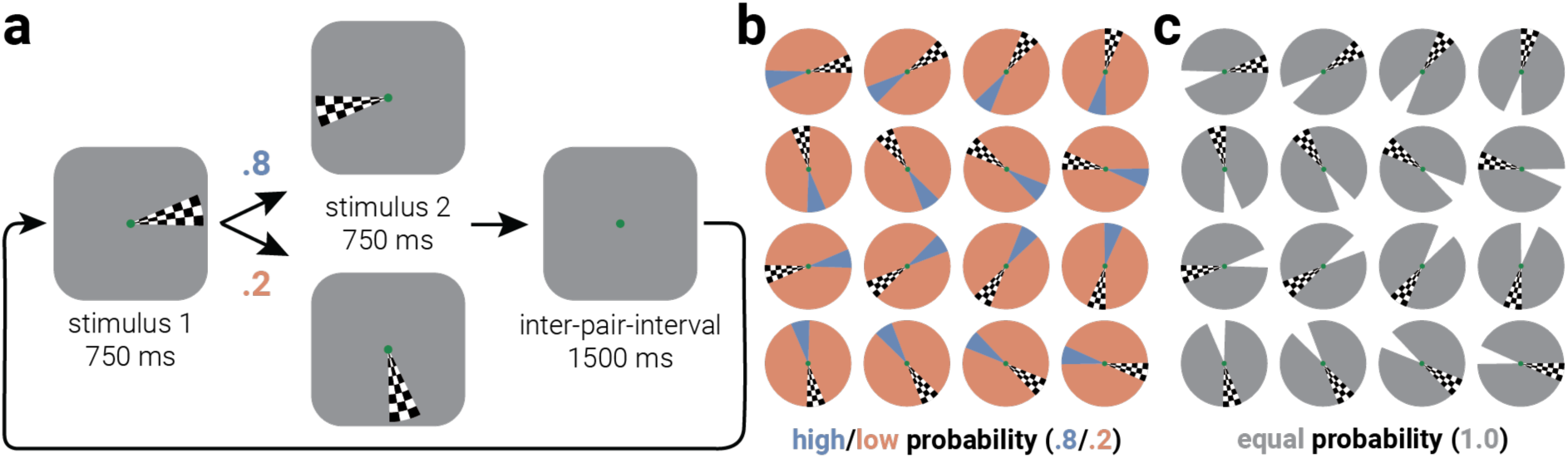
Manipulating probability within a population receptive field mapping design. **a**) Illustration of experimental design. A checkerboard wedge was presented at one of 16 possible polar angles, followed by a second wedge presented either at the location 180° from the first wedge (80% probability) or at any of the other (14) locations, excluding the initial and opposite locations (20% probability). This paired sequence was repeated, with a uniform distribution of initial wedge locations, throughout the block, separated by a blank inter-pair-interval. Participants were tasked with detecting brief and infrequent contrast reductions of the wedge stimuli. **b**) Illustration of the 16 possible wedge locations and the locations at which subsequent wedges could be presented. **c**) As a baseline to compare against high- and low-probability presentations, we also included blocks in which the second wedge was always presented with equal probability (“random” stimuli).

As a baseline, to determine directionality of change associated with expectation, a single block in each experimental session (either the 3^rd^ or 4^th^ block) comprised trials in which the second wedge was presented at any of the 16 possible locations, excluding the same location of the first and its polar opposite (equal probability, “random” stimuli; **Fig. 1c**). To maintain engagement, throughout the experimental sessions, participants were tasked with pressing a button when they detected wedge contrast decrements (−25% contrast). Contrast decrements occurred infrequently (20% of trials) for the duration of one of the polarity alternations (150 ms) on either the first or second wedge.

### Magnetic resonance imaging and data preprocessing

Magnetic resonance scanning was conducted on a Magnetom 7T research whole-body MRI scanner (Siemens Healthcare, Erlangen, Germany) equipped with a 32-channel head coil (Nova Medical, USA). Blood-oxygen level-dependent (BOLD) fMRI data were acquired with a gradient echo echo-planar imaging (EPI) sequence for experimental [echo time (TE), 20 ms; repetition time (TR), 750 ms; voxel size, 2 mm isotropic, 30 slices covering occipital cortex] and retinotopic scans [TE, 24 ms; TR, 1000 ms; voxel size, 1.2 mm isotropic, 36 slices covering occipital cortex]. Slice orientation was close to coronal but rotated about the mediolateral axis with the ventral edge of the volume more anterior to ensure coverage of lateral occipital cortex. A T1-weighted, 3D MP2-RAGE pulse sequence, anatomical scan (voxel size, 0.8 mm isotropic) was also acquired for each participant. These anatomical scans were used to register the functional data across scanning sessions, restrict the analysis of BOLD activity to grey matter voxels, and create flattened surfaces on which cortical activity was visualized.

Anatomical scans for each participant were aligned to the anterior commissure - posterior commissure (AC-PC) line, and inflated and flattened surfaces were rendered with BrainVoyager QX (BrainInnovation, Maastricht, The Netherlands). Functional data were preprocessed with slice timing correction, head motion correction, and high-pass filtering, before being aligned to the participant’s anatomical scan and aligned to the AC-PC line.

### Population receptive field mapping

The fMRI data from the experimental blocks were used to obtain independent estimates of the pRFs using a custom MATLAB toolbox for pRF analysis (SamSrf v9.9, https://github.com/samsrf). We predicted the neural response of each vertex in the cortical surface mesh from the overlap of a binary aperture describing the position of the wedge stimuli within each scanning volume with a model of the underlying pRF. We then convolved this with a canonical haemodynamic response function (HRF) to predict the BOLD signal in the experimental sessions. The binary aperture was a two-dimensional mask (100 × 100) corresponding to the stimulus location on the screen at each time point. We estimated a unidimensional pRF model of polar angle using wedge apertures (Dumoulin & Wandell, 2008), constrained to locations on a circle around fixation with radius 6° (half the outer eccentricity of the wedge stimulus). This effectively produced a one-dimensional tuning model defined by two parameters: μ indicated the preferred polar angle of the voxel and σ corresponded to the tuning width. To compare the width of pRFs in response to the different stimuli (first stimulus and high/low/equal probability second stimulus), which were interleaved throughout the experimental blocks, we modified the standard pRF model to include three separate σ parameters to describe tuning width, σ_1st_ for the initial stimulus in each pair, σ_Low_ for the stimuli that violated expectations, and σ_High_ for expected stimuli (i.e., occurring with different probabilities). This modification, combined with the large data volume produced by each fMRI block, increased the computing power required to fit parameters. This precluded the use of parameter optimization. We therefore only used an extensive grid search to determine the pRF parameters that yielded the best fit of the predicted time series with the observed data. For the same reason, we performed the analysis separately on data from each experimental block, before averaging the resultant parameter estimates.

While performing the pRF mapping analysis, we found that the number of events included in the analysis influenced the σ estimate, such that more events produced more narrowly tuned pRF estimates. This was particularly evident for the first stimulus, which occurred more frequently than the high/low/equal probability stimuli, which were split between the second stimulus presentations. For the purposes of our research question, the difference between the first and second stimulus was unimportant. However, the high, low, and equal probability stimuli were also presented with different frequencies. Specifically, although there were 240 first events and 240 second events, of the second events in the main experimental blocks, there were 192 high probability stimuli but only 48 low probability stimuli. The pRF mapping analysis was run separately on data from each block, so to match the number of events included in the pRF analysis for high and low probability conditions, we labelled 144 of the high probability stimuli as first events (388 first events, 48 high probability events, and 48 low probability events). All of the second events in the baseline blocks were equal probability stimuli. Thus, to match the analysis of these events to those in the main blocks, we relabelled 144 of the second events as first events, 48 as equal probability (A), and 48 as equal probability (B; 388 first events, 48 equal probability A events, and 48 equal probability B events). The parameter estimates of A and B events were then averaged to produce a single estimate for equal probability events.

### Regression analysis

As noted above, we developed a modified pRF mapping analysis to compare the width of receptive fields between stimulus probabilities. In addition to performing a validation using simulated data to confirm that the method produced sensible parameter estimates when the ground truth was known, we also confirmed the results using an alternative analytical approach. In this approach, we z-normalized the time series of each voxel within an experimental block, before temporally concatenating across experimental blocks. We then created a binary design matrix comprising the angular position and stimulus probability of each wedge in the series, and convolved this with a canonical HRF. Combining the design matrix with the normalized time series data from each voxel, we used regression to compute β weights of each position-probability combination. We then used a least-squares procedure to fit a von Mises function to the position weights separately within each probability group, with μ indicating the preferred polar angle of the voxel and κ corresponding to the concentration of the β weights.

### Retinotopy

In a separate session, we localized regions of interest (ROIs) for each participant using standard retinotopic mapping procedures. Retinotopically organized visual areas V1, V2, V3, V3AB, and V4 were defined with polar and eccentricity maps, which were generated by BOLD responses to rotating wedge and expanding concentric ring stimulus presentations, respectively (DeYoe, Carman, Bandettini, Glickman, Wieser, et al., 1996; M. Sereno et al., 1995). Analysis of retinotopic data was performed using the Fourier transformation-based method, as documented in standard retinotopy studies (DeYoe, Carman, Bandettini, Glickman, Wijnen, et al., 1996; Engel et al., 1994; M. I. Sereno et al., 1995).

### EEG recording and preprocessing

The EEG experiment was conducted in a dark, sound-proof, and electromagnetically shielded room. Stimuli were presented on a display monitor (ASUS VG248QE 144 Hz 24’’ 1920×1080 HDMI Gaming Monitor) via MATLAB R2020a (The MathWorks, Inc., Matick, MA) and PsychToolbox (v3.0, http://psychtoolbox.org/; Pelli, 1997). Participants were instructed to place their head on a chin-forehead rest, which controlled the viewing distance at 70 cm with the screen subtending 41.47° x 23.33° of visual angle.

We employed 64-channel active Ag/AgCl electrodes with the arrangement on the scalp aligning with the 10–20 international standard (Oostenveld & Praamstra, 2001), via a nylon cap and an online reference point at FCz. EEG data were captured at a sampling rate of 1024 Hz using a 24-bit A/D conversion and recorded on a BrainVision ActiCap system (Brain Products GmbH, Gilching, Germany). The EEG data pre-processing was conducted using EEGLAB version 2021.1 (Delorme & Makeig, 2004). The data sampling rate was initially reduced to 256 Hz. To eliminate slow baseline fluctuations, a 0.1 Hz high-pass filter was applied, and a 45.0 Hz low-pass filter was employed to filter out high-frequency noise. Next, the *clean_artifacts* function was applied, without high-pass filtering, and the most lenient output (BUR) was used. Following this, independent component analysis (ICA) was performed using the automatic SASICA pipeline (Chaumon et al., 2015) to identify and remove components corresponding to ocular and other artifacts, including eye blinks. The final data were epoched from -200 ms to 500 ms relative to the onsets of the wedge stimuli.

### EEG decoding

To characterise neural representations of the visual stimuli, we used an inverted modelling approach to reconstruct the location of the wedges from the EEG data (Brouwer & Heeger, 2011). A theoretical (forward) model was chosen that described the measured activity in the EEG sensors given the location of the presented wedge. The forward model was then used to obtain the inverse model that described the transformation from EEG sensor activity to stimulus location (Lee & Rideaux, 2025; Rideaux, 2024). The forward and inverse models were based on neural responses to the first wedge within each pair to obtain the inverse model on which responses to the second wedge were decoded.

Similar to previous work (Brouwer & Heeger, 2009), the forward model comprised five hypothetical channels, with evenly distributed idealized location preferences between 0° and 360°. Each channel consisted of a half-wave rectified sinusoid raised to the fifth power. The channels were arranged such that a tuning curve of any location preference could be expressed as a weighted sum of the five channels. The observed EEG activity for each presentation could be described by the following linear model:

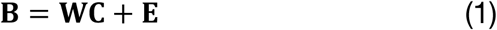

where **B** indicates the (*m* sensors × *n* presentations) EEG data, **W** is a weight matrix (*m* sensors × 5 channels) that describes the transformation from EEG activity to stimulus location, **C** denotes the hypothesized channel activities (5 channels × *n* presentations), and **E** indicates the residual errors.

To compute the inverse model, we estimated the weights that, when applied to the data, reconstructed the underlying channel activities with the least error. In line with previous magnetencephalography work (Kok et al., 2017; Mostert et al., 2015), when computing the inverse model, we deviated from the forward model proposed by (Brouwer & Heeger, 2009) by taking the noise covariance into account to optimize it for EEG data, given the high correlations between neighbouring sensors. We then estimated the weights that, when applied to the data, reconstructed the underlying channel activities with the least error. Specifically, **B** and **C** were demeaned such that their average over presentations equalled zero for each sensor and channel, respectively. The inverse model was then estimated from the responses to the first wedge in each pair. The hypothetical responses of each of the five channels were calculated from the training data, resulting in the response row vector **c***_train_*_,*i*_ of length *n*_train_ presentations for each channel *i*. The weights on the sensors **w***_i_* were then obtained through least squares estimation for each channel:

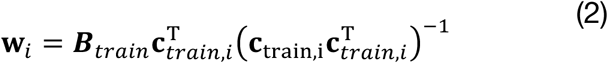

where ***B****_train_* indicates the (*m* sensors × *n*_train_ presentations) training EEG data. Subsequently, the optimal spatial filter **v***_i_* to recover the activity of the *i*th channel was obtained as follows (Mostert et al., 2015):

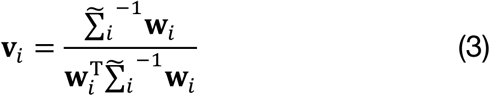

where Σ̃_*i*_ is the regularized covariance matrix for channel *i*. Incorporating the noise covariance in the filter estimation leads to suppression of noise that arises from correlations between sensors. The noise covariance was estimated as follows:

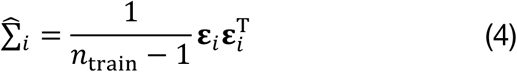

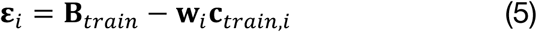

where *n*_train_ is the number of training presentations. For optimal noise suppression, we improved this estimation by means of regularization by shrinkage using the analytically determined optimal shrinkage parameter (Mostert et al., 2015), yielding the regularized covariance matrix Σ̃_*i*_.

For each presentation, we decoded location by converting the channel responses to polar form:

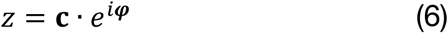

and calculating the estimated angle:

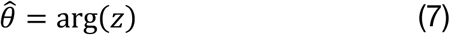

where **c** is a vector of channel responses and ***φ*** is the vector of angles at which the channels peak. As outlined above for the analysis of fMRI data, we balanced the number of trials included across conditions (low probability, high probability, equal probability) to eliminate any potential confounding influence of unbalanced observations.

From the decoded locations, we computed three estimates: *accuracy, precision*, and *bias* (Harrison et al., 2023; Rideaux, Bays, et al., 2025; Rideaux et al., 2023). Accuracy represents the similarity of the decoded location to the presented location (Kok et al., 2017), and was expressed by projecting the mean resultant (averaged across presentations within the same stimulus location) of the difference between decoded and wedge location onto a vector with 0°:

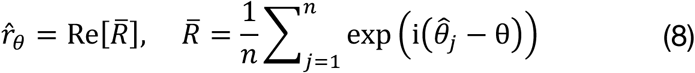

Precision was estimated by calculating the angular deviation (Zar, 1999) of the decoded locations for each of the 16 unique locations:

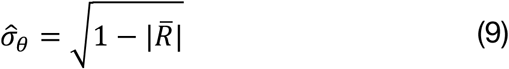

and normalized, such that values ranged from 0 to 1, where 0 indicates a uniform distribution of decoded location across all locations (i.e., chance-level decoding) and 1 represents perfect consensus among decoded locations:

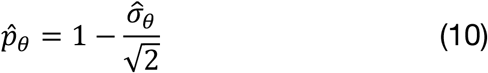

Bias was estimated by computing the circular mean of the angular difference between decoded and presented location:

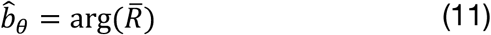

We included only the parietal, parietal-occipital, and occipital sensors (20 in total) in the analyses to *i*) reduce the influence of signals produced by any eye movements and blinks, *ii*) for consistency with previous decoding studies (Buhmann et al., 2024; Rideaux, 2024; Rideaux, Bays, et al., 2025), and *iii*) to improve decoding accuracy by restricting sensors to those that carried spatial position information.

### Statistical analyses

Statistical analyses were performed in MATLAB v2020a and CircStat Toolbox v1.12.0.0 (Berens, 2009). For pRF mapping analyses of fMRI data, we used only the fitted parameters of those vertices that had a goodness of fit, R^2^ higher than 0.1. For both pRF and regression analyses of fMRI data, two-way repeated measures analysis of variance was used to test for main effects in width parameter estimates for ROI and stimulus probability (high, low, and equal). For analyses of EEG estimates as a function of time, a cluster correction was applied to remove spurious significant differences. First, at each time point, the main effect size of stimulus probability for ERP amplitude/accuracy/precision was calculated using a repeated measure analysis of variance. Next, we calculated the summed value of these statistics (separately for positive and negative values) within contiguous temporal clusters of significant values. We then simulated the null distribution of the maximum summed cluster values using permutation (*n*=5000) of the probability labels, from which we derived the 95% percentile threshold value. Clusters identified in the data with a summed effect-size value less than the threshold were considered spurious and removed.

## RESULTS

For the fMRI data, we used a modified pRF mapping approach to compute the preferred polar angle and spatial tuning width associated of neural populations in visual cortex with: *i*) the first stimulus in the paired presentation, and the second stimulus when it was probabilistically *ii*) high, iii) low, or iv) equal. **Figure 2** shows the results of the analysis projected onto a flattened cortical surface for a single participant. We found that the correspondence between the model and the data (R^2^) was highest around retinotopically defined early visual areas (V1, V2, V3, V3AB, & V4; **Fig. 2a**). The correspondence between the preferred polar angle derived from pRF mapping and the phase-encoded retinotopy was high (average Pearson’s r=0.95 across participants; **Fig. 2b**). Similar to the model-fit results, pRF polar angle width was sharpest in early visual cortex; **Figure 2c** shows the width of the response to the first stimulus in the paired presentation. Critically, to assess the influence of probability on spatial tuning, we were concerned with the difference between pRF width in response to high and low probability stimuli (**Fig. 2d**).

**Figure 2.**
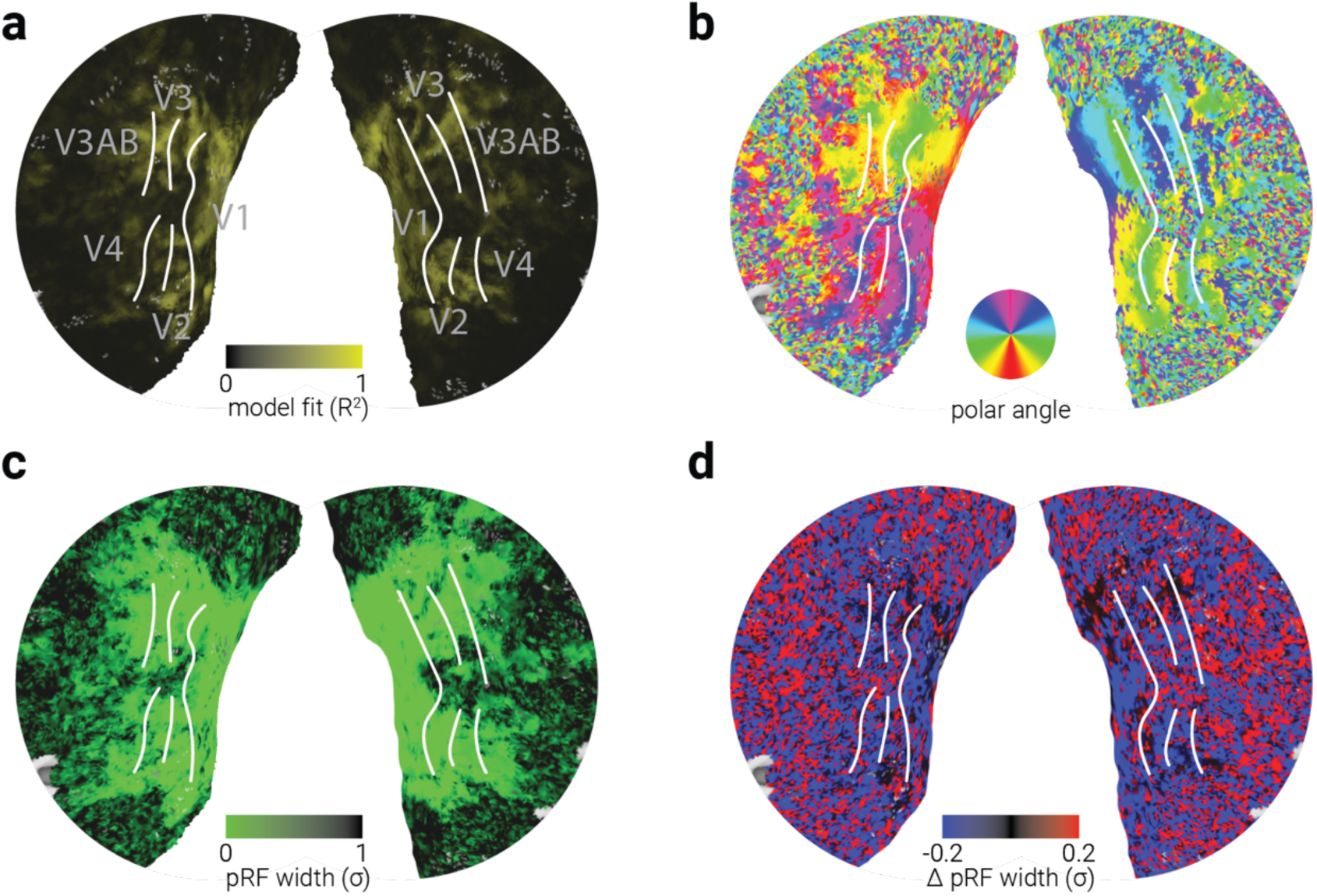
Population receptive field properties across the cortical sheet. The flattened cortical surface of a representative subject (S1) overlayed with the following parameters derived from the pRF mapping analysis: (**a**) model fit, (**b**) preferred polar angle, (**c**) pRF width in response to the first wedge stimulus, and (**d**) the difference in pRF width in response to high and low probability wedges (blue values indicate sharper tuning for high probability stimuli). White lines demarcate the borders of visual areas, defined using phase-encoded analysis of a standard retinotopic fMRI design collected in a separate session.

To test the influence of probability on pRF width across the visual cortex, we calculated the average pRF width of voxels in each ROI, for high, low and equal probability stimuli. There were significant main effects of ROI (*F*_4,36_=3.78, *p*=.012) and stimulus probability (*F*_2,18_=22.70, *p*=1.20e^-5^), and a significant interaction between the two (*F*_8,72_=6.58, *p*=2.04e^-6^), indicating that probability influenced the width of pRFs more in early than later visual areas. Comparison of high and low probability pRF widths with those of equal probability indicated that spatial tuning was significantly narrower for stimuli presented at high probability locations, whereas no change was observed for stimuli at low probability locations (**Fig. 3a**). The results from an additional regression analysis were broadly consistent with those from the pRF mapping: there were significant main effects of both ROI (*F*_4,36_=16.06, *p*=1.24e^-7^) and stimulus probability (*F*_2,18_=7.21, *p*=.005), but the interaction was not significant (*F*_8,72_=1.68, *p*=.117). While less clear than the pRF mapping results, the regression analysis results also indicated that the effect of probability on the fidelity of neural representations was driven by sharpened tuning in response to high probability stimuli (**Fig. 3b**).

**Figure 3.**
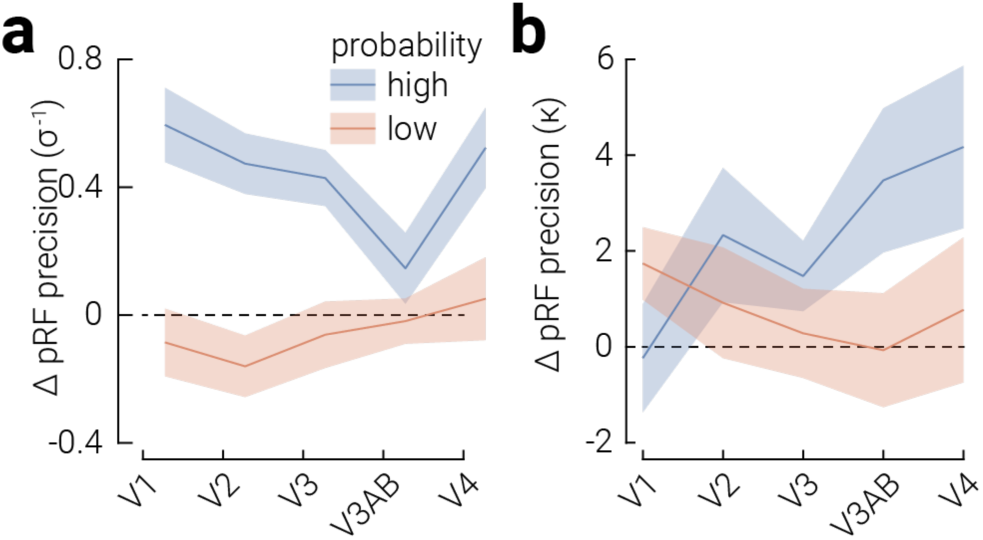
Sharper population receptive field tuning for high probability wedges. **a**) The difference in pRF precision relative to equal probability stimuli for high and low probability stimuli, as a function of visual area, as established from the pRF mapping analysis. **b**) Same as (**a**) but derived from an alternative regression analysis (see Methods section for details). Shaded regions indicate ±SEM.

The fMRI results support the sharpening account of predictive processing, consistent with previous work (Kok et al., 2012). However, recent theoretical accounts of predictive processing have proposed that an initial period of sharpening (i.e., pre-activation) may be followed by a later period of dampening in response to expected stimuli (Press et al., 2020). The proposed temporal separation of these putative effects exceed the temporal resolution of BOLD fMRI, so if both sharpening and dampening were occurring in the current paradigm, we would be unable to distinguish them with fMRI alone. Further, in our experimental design, stimuli that were probabilistically likely were also presented at larger angular offsets from the preceding stimulus than those that were unlikely or equally probable. Thus, it is possible that local spatiotemporal stimulus properties may have contributed to the differences observed between probability conditions. That is, differences between conditions may have been due to the distance between wedge pairs, rather than expectation. Testing the influence of local spatiotemporal stimulus properties on tuning width would require estimating tuning properties at relative angular offsets from the first stimulus. This is theoretically possible to perform using the modified pRF mapping approach developed here to separately assess the responses associated with the three probability conditions. However, in practice, these methods are ill suited to this kind of analysis, as they require responses to a full set of polar locations to estimate tuning parameters; splitting the data according to angular offset would reduce the reliability of the results considerably. A better approach is to use multivariate decoding, which can estimate precision based on the response at a single location, rather than a full set of locations. We therefore performed the same experiment while recording neural activity with EEG. Further, to assess whether spatiotemporal stimulus properties influenced the results observed in the fMRI experiment, we used multivariate decoding to assess representational fidelity at each angular offset between the first and second stimulus.

We began by comparing the aggregate (univariate) EEG responses to stimuli of different probabilities. Analysis of the univariate response to high, low, and equal probability wedge stimuli revealed two periods of significant difference between the three stimulus probabilities, around the P2 and N3 evoked components (**Fig. 4a**). Similar to the fMRI results, differences in univariate amplitude between the probabilities were driven by responses to high probability stimuli, which were larger than those in response to low and equal probability stimuli. Note, however, that this result runs counter to *expectation suppression*, which predicts a *reduced* response to expected stimuli (Kok et al., 2012; Teufel et al., 2018; Yon et al., 2018).

**Figure 4.**
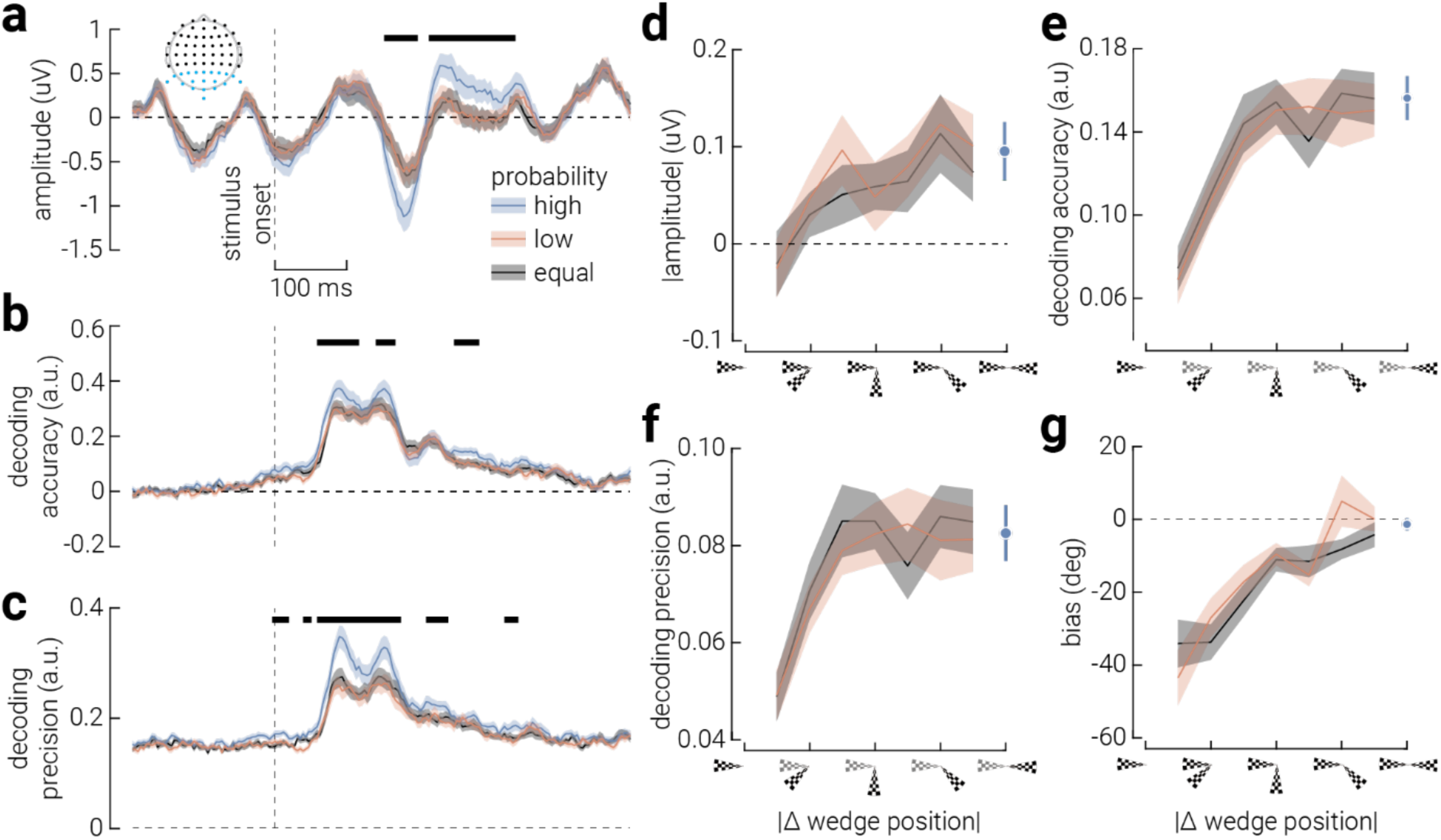
Differences in neural tuning due to spatiotemporal properties, not probability. **a**) Univariate amplitude of evoked responses to high, low, and equal probability stimuli between -200 and 500 ms around stimulus onset. **b, c**) The same as (**a**), but for decoding (**b**) accuracy and (**c**) precision. Note, the uniformly elevated decoding precision is due to the anisotropic neural representation of polar angle, which biases the likelihood of the decoder predicting some locations over others, even in the absence of target related signals. Horizontal black bars indicate cluster corrected periods of significant main effects of stimulus probability. The inset in (**a**) indicates the sensors included across all analyses (blue dots). **d**) Absolute univariate amplitude of evoked responses to high, low, and equal probability stimuli, as a function of the absolute angular distance from the first stimulus. **e**-**g**) Same as (**d**), but for decoding (**e**) accuracy, (**f**) precision, and (**g**) bias. Shaded regions and error bars indicate ±SEM. The absence of data at zero delta position is because the second wedge was never presented at the same location as the first. Similarly, the high probability wedges were always presented 180° from the preceding wedge, while low and equal probability wedges were never presented at this offset.

We next used inverted encoding modelling to decode the location of wedge stimuli from the multivariate pattern of activity across EEG sensors, and from this we estimated decoding *accuracy*, *precision*, and *bias*. There were three periods in which decoding accuracy was significantly different between high, low, and equal probability stimuli (**Fig. 4b**). Differences in decoding accuracy emerged early in the epoch (∼60 ms), during the initial surge in decoding accuracy associated with feedforward activity, and largely occurred in periods that were distinct from those for later univariate responses (∼150 ms). This is consistent with previous work showing that inverted encoding can reveal phenomena obscured in the univariate signal (Buhmann et al., 2024). A similar pattern of results was apparent for decoding precision (**Fig. 4c**). We also found that decoding precision was uniformly above-chance, even before stimulus onset. This indicates a non-uniform probability of predicting certain locations over others, which likely reflects the anisotropic representation of polar angle in early visual cortex (Freeman et al., 2013; Sasaki et al., 2006). There were some periods over which there were significant differences in precision that we did not observe for accuracy, including just after stimulus onset but prior to cortical stimulus processing (0-40 ms). Similar non-significant differences were apparent for decoding accuracy, suggesting that precision may be a more sensitive measure for this effect. Given the early timing of the effect, this difference may reflect anticipatory activity (Kok et al., 2017).

Overall, findings from the EEG decoding analyses were broadly consistent with those from the fMRI experiment: the location of high probability wedges was decoded more accurately and precisely, indicating sharper tuning to these stimuli. However, as mentioned previously, the angular offset of the high probability stimuli from the first stimulus was larger than that for the low and equal probability stimuli. To assess the influence of these local spatiotemporal differences we compared responses to low and equal probability stimuli, because in these conditions the local spatiotemporal properties were matched, while probability differed. In particular, in the low probability condition, stimuli were most likely to be presented in the polar opposite location to the first stimulus, whereas in the equal probability condition stimuli were never presented at this location.

If wedges were expected at the polar opposite location, and this caused sharpening of responses, we might expect to observe a similar, but reduced, effect in response to nearby low probability stimuli, but not to nearby equal probability stimuli (where this was the least likely location). We tested for this in the EEG data by averaging results across time (50–500 ms) and splitting trials according to the absolute angular distance between the first and second wedges within each pair. For univariate responses, given the bidirectional nature of differences observed over time, i.e., both positive and negative differences between conditions, we calculated the absolute difference in amplitude evoked by the stimuli. Responses to low probability stimuli increased as a function of angular distance, reaching the same amplitude as for high probability stimuli at the smallest offset (**Fig. 4d**). This result is consistent with sharper tuning for expected stimuli. However, the same pattern was apparent for the equal probability stimuli, for which the opposite location had zero probability. Indeed, we observed the same pattern of results for both decoding accuracy and precision (**Fig. 4e**, **f**), which co-occurred with a repulsive bias away from the location of the first stimulus (**Fig. 4g**).

Recall that the equal probability blocks were interleaved with the main experimental blocks. Thus, a possible explanation for the lack of difference between the low and equal probability stimuli is that participants’ expectations developed in the low probability blocks were maintained in the equal probability blocks. To test this, we separately analysed data from the first and last third of the equal probability blocks. If participants’ expectations about the likely location of the second wedge carried over from the preceding blocks, any such effect should be degraded over the course of the equal probability block. However, there was no difference in decoding accuracy between the start and end of the equal probability block (**Supplementary Fig. 1**). Similarly, there was no difference in task performance between the three stimulus probabilities in either the fMRI experiment (average overall performance, 66.3%; *F*_2,18_=0.78, *p*=.472) or the EEG experiment (average, 68.0%; *F*_2,46_=1.94, *p*=.155) indicating that the effects were not a result of differential task engagement.

## DISCUSSION

The temporal context in which stimuli are embedded can drive expectations that support adaptive behaviours and efficient sensory processing. Here we combined pRF mapping of fMRI BOLD responses with inverted encoding of EEG recordings to adjudicate between sharpening and dampening accounts of how expectation influences representational fidelity of simple visual stimuli. The fMRI results appeared to support a sharpening account of expectation (Kok et al., 2012; Teufel et al., 2018; Yon et al., 2018); however, analysis of EEG data revealed that differences observed between processing of high and low probability stimuli were unlikely to reflect expectation established by our probability manipulation. Instead, the findings were more parsimoniously explained by local spatiotemporal properties of the stimuli.

We manipulated expectancy by presenting stimuli at locations with either high (80%), low (20%), or equal probability. The results from both the fMRI and EEG experiments provided converging evidence that high probability stimuli elicited sharper spatial representations. However, the univariate responses to these stimuli were also higher, directly contradicting expectation suppression (Summerfield et al., 2008), which is thought to conserve metabolic expenditure in response to expected stimuli. If the differences in spatial tuning were caused by the probability manipulation, two patterns of results should be hypothesised for low and equal probability stimuli as a function of angular distance from the high probability location. First, if expectation tuning is highly precise, high probability stimuli should be more accurately decoded, and there should be no effect of angular distance for either low or equal probability stimuli, because only stimuli presented exactly at the expected location should elicit an anticipatory effect. Alternatively, if expectation tuning is not highly precise, then low probability stimuli (but not equal probability stimuli) presented near the expected location should be decoded more accurately, because stimuli presented at near-to-expected locations should also elicit an anticipatory effect. However, neither of these hypotheses was supported. Instead, decoding accuracy and precision for low and equal probability stimuli were the same as for high probability stimuli up to offsets of ∼45°, at which point they both declined as a function of angular distance.

These results suggest that differences in spatial tuning were caused by local spatiotemporal stimulus properties, specifically, the angular distance between the first and second wedges in each pair. This would explain why there was no difference between low and equal probability stimuli, and why differential decoding accuracy and precision were highest around the location of the first wedge and lowest around the (high probability) opposite location. The changes in decoding accuracy and precision were accompanied by a repulsive bias away from the initial wedge location, which is consistent with a sensory adaptation effect (Whitaker et al., 1997). However, the presentation duration of the first stimulus (750 ms) was considerably shorter than the standard duration used to induce sensory adaptation (30 s; Clifford, 2002), and other aspects of the phenomenon we observed here seem inconsistent with such a process. For instance, previous EEG decoding work has shown that adaptation to orientation induces both increased univariate responses and decoding precision around the adapted location (Rideaux et al., 2023), whereas here we observed the opposite.

Recent work on the influence of temporal context on sensory processing has shown that repeated events are remembered less precisely than those that change (Lee & Rideaux, 2025). This could be explained by an underlying expectation that sensory events will stay the same, or repeat, rather than change. The notion of an expectation for visual stability may seem counterintuitive given that retinal input is often changing, e.g., due to saccades, self-motion, and world-motion. However, for a prediction to be useful (e.g., to reduce metabolic expenditure or speed up responses) it must be somewhat precise. A prediction that retinal input will change is not necessarily useful unless it can specify what it will change to. If change is predicted, given retinal input of *x_t_* at time *t*, the range of possible values of *x_t_*_+1_ is infinite. By contrast, if no change is predicted, such that *x_t_*_+1_=*x_t_*, then a precise prediction can be established and utilized. There is, of course, other information that can be used to reduce the parameter space of predicted change from *x_t_*, for example, the value of *x_t_*_-1_. However, these sources of expectation are not mutually exclusive. Indeed, across the infinite distribution of changes from *x_t_*, zero change will occur more frequently than any other value, so it seems plausible that the brain would be sensitive to this pattern.

If there is indeed an expectation of temporal stability, the current findings would support the dampening account of expectation. Expected stimuli (those nearby to the repeat location) were suppressed both at the univariate level, due to expectation suppression, and their fidelity was reduced, as indicated by both broader pRFs and reduced decoding accuracy and precision. Moreover, the shape of the differential decoding as a function of angular distance suggests suppression around the expected (repeat) location, rather than enhancement around the least expected (opposite) location. Ecologically, suppressing expected sensory inputs may be adaptive because these events are already captured by the system’s predictive model, whereas unexpected inputs must be encoded precisely in order to effectively update the model (Soltani & Izquierdo, 2019).

Taken together, the current findings are consistent with previous work on predictive processing, suggesting that expected events are suppressed, and that unexpected events are more reliably decoded from neural activity (Richter et al., 2022; Tang et al., 2018) and behaviourally reproduced more precisely (Rideaux, Dang, et al., 2025). In contrast, they seem to contradict studies that have shown expected events are decoded more reliably (Kok et al., 2012) and produce more precise behavioural responses (Rohenkohl et al., 2014). The interaction between global probability and local spatiotemporal properties observed in the current study provides an example of the challenging nature of isolating the influence of expectation and may offer insight into the discrepant findings associated with this phenomenon. The fact that the probability manipulation did not induce detectable neural effects is also consistent with a growing body of evidence indicating that while probabilistic cuing paradigms produce robust behavioural effects, they do not always elicit neural responses that can be reliably detected with human neuroimaging methods (den Ouden et al., 2023, 2025; Hu et al., 2025).

Decoding accuracy is frequently used as a proxy for tuning width, as more sharply tuned responses are more distinguishable from responses to other points along a feature dimension and will thus result in more accurate decoding. However, the same outcome could also be produced, without changing tuning width, by increasing the temporal reliability of the response (Stehr et al., 2023). By applying pRF mapping and inverted decoding to neural recordings made during the same experimental protocol, we have provided the first validation across two imaging methods (fMRI and EEG) that decoding accuracy does indeed reflect narrower tuning. Interestingly, decoding precision, which is seldom computed in neural decoding work, appeared to be a more sensitive measure of the experimentally induced changes in neural activity. This is consistent with previous work suggesting that decoding precision may be more sensitive to task demands than accuracy (Harrison et al., 2023).

A clear limitation of the current study is that the global expectancy manipulation did not appear to be effective. While we found evidence of dampening, given the partial alignment of the pattern of results with sensory adaptation, further work is needed to determine whether the neural phenomena observed here were due to expectation of stability or adaptation. Another limitation of the study is that we did not employ a behavioural marker of expectation, which could have been used to cross-validate the efficacy of the global spatiotemporal manipulation. Such a behavioural marker could further have been used to compare performance between spatiotemporally proximal and distal pairs, to adjudicate between adaptation and expectation accounts. However, previous behavioural work has shown a striking similarity between the effects of local and global spatiotemporal statistical manipulations on response time, accuracy, and precision of responses (Lee & Rideaux, 2025), suggesting the neural effects observed here are consistent with expectation of stability.

In summary, our findings highlight both the promise and the difficulty of disentangling expectation from other sources of temporal context in sensory processing. While a superficial interpretation of the results implied sharpened tuning for expected stimuli, additional analyses of the EEG data suggest instead that local spatiotemporal properties provide a more parsimonious explanation; however, more work is needed to rule out adaptation. Viewed through the lens of predictive processing, the current results align more closely with a dampening account, whereby expected inputs are suppressed and unexpected inputs are represented with greater fidelity. Beyond their theoretical implications, our findings underscore the importance of approaches which combine complementary imaging methods and include refined decoding metrics for advancing our understanding of how the brain leverages temporal context to guide perception.

## Acknowledgements

RR was supported by an Australian Research Council (ARC) Discovery Early Career Researcher Award (DE210100790) and a National Health and Medical Research Council (NHMRC; Australia) Investigator Grant (2026318). JBM was supported by an NHMRC Investigator Grant (2010141).

## SUPPLEMENTARY MATERIAL

**Figure S1.**
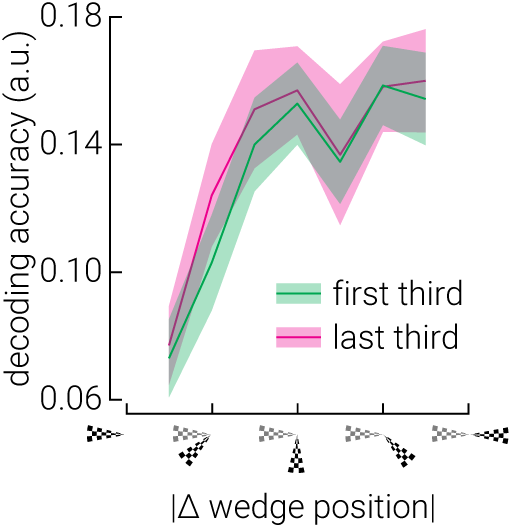
No evidence for change in expectation across the equal probability blocks. Decoding accuracy of equal probability stimuli from the first and last third of trials in the block, as a function of the absolute angular distance from the first stimulus. Shaded regions indicate ±SEM.

## REFERENCES

1. Barlow, H. B. (1961). Possible Principles Underlying the Transformations of Sensory Messages. In W. A. Rosenblith (Ed.), Sensory Communication (pp. 216–234). The MIT Press. 10.7551/mitpress/9780262518420.003.0013

2. Berens, P. (2009). CircStat: A MATLAB Toolbox for Circular Statistics. Journal of Statistical Software, 31(10). 10.18637/jss.v031.i10

3. Brouwer, G. J., & Heeger, D. J. (2009). Decoding and reconstructing color from responses in human visual cortex. Journal of Neuroscience, 29(44), 13992–14003. 10.1523/JNEUROSCI.3577-09.2009

4. Brouwer, G. J., & Heeger, D. J. (2011). Cross-orientation suppression in human visual cortex. Journal of Neurophysiology, 106(5), 2108–2119. 10.1152/jn.00540.2011

5. Buhmann, Z., Robinson, A. K., Mattingley, J. B., & Rideaux, R. (2024). Inverted encoding of neural responses to audiovisual stimuli reveals super-additive multisensory enhancement. eLife, 13. 10.7554/eLife.97230.1

6. Chaumon, M., Bishop, D. V. M., & Busch, N. A. (2015). A practical guide to the selection of independent components of the electroencephalogram for artifact correction. Journal of Neuroscience Methods, 250, 47–63. 10.1016/j.jneumeth.2015.02.025

7. Clifford, C. W. G. (2002). Perceptual adaptation: Motion parallels orientation. Trends in Cognitive Sciences, 6(3), 136–143. 10.1016/S1364-6613(00)01856-8

8. Delorme, A., & Makeig, S. (2004). EEGLAB: An open source toolbox for analysis of single-trial EEG dynamics including independent component analysis. Journal of Neuroscience Methods, 134(1), 9–21. 10.1016/j.jneumeth.2003.10.009

9. den Ouden, C., Kashyap, M., Kikkawa, M., & Feuerriegel, D. (2025). Limited Evidence for Probabilistic Cueing Effects on Grating-Evoked Event-Related Potentials and Orientation Decoding Performance. Psychophysiology, 62(5), e70076. 10.1111/psyp.70076

10. den Ouden, C., Zhou, A., Mepani, V., Kovács, G., Vogels, R., & Feuerriegel, D. (2023). Stimulus expectations do not modulate visual event-related potentials in probabilistic cueing designs. NeuroImage, 280, 120347. 10.1016/j.neuroimage.2023.120347

11. DeYoe, E. A., Carman, G. J., Bandettini, P. A., Glickman, S., Wijnen, J. P., Cox, R., Miller, D., & Neitz, J. (1996). Mapping striate and extrastriate visual areas in human cerebral cortex. Proceedings of the National Academy of Sciences of the United States of America, 93(6), 2382–2386. 10.1073/pnas.93.6.2382

12. DeYoe, E. A., Carman, G. J., Bandettini, P., Glickman, S., Wieser, J., Cox, R., Miller, D., & Neitz, J. (1996). Mapping striate and extrastriate visual areas in human cerebral cortex. Proceedings of the National Academy of Sciences, 93(6), 2382–2386. 10.1073/pnas.93.6.2382

13. Dumoulin, S. O., & Wandell, B. A. (2008). Population receptive field estimates in human visual cortex. NeuroImage, 39(2), 647–660. 10.1016/j.neuroimage.2007.09.034

14. Engel, S. A., Rumelhart, D. E., Wandell, B. A., Lee, A. T., Glover, G. H., Chichilnisky, E. J., & Shadlen, M. N. (1994). FMRI of human visual cortex [5]. Nature, 369(6481), 525. 10.1038/369525a0

15. Feuerriegel, D. (2024). Adaptation in the visual system: Networked fatigue or suppressed prediction error signalling? Cortex, 177, 302–320. 10.1016/j.cortex.2024.06.003

16. Freeman, J., Heeger, D. J., & Merriam, E. P. (2013). Coarse-Scale Biases for Spirals and Orientation in Human Visual Cortex. Journal of Neuroscience, 33(50), 19695–19703. 10.1523/JNEUROSCI.0889-13.2013

17. Friston, K. (2005). A theory of cortical responses. Philosophical Transactions of the Royal Society B: Biological Sciences, 360(1456), 815–836. 10.1098/rstb.2005.1622

18. Harrison, W. J., Bays, P. M., & Rideaux, R. (2023). Neural tuning instantiates prior expectations in the human visual system. Nature Communications, 14(1), Article 1. 10.1038/s41467-023-41027-w

19. Hu, Z., Tran, D. M. D., & Rideaux, R. (2025). Multimodal evidence challenges the effectiveness of probabilistic cueing for establishing sensory expectations (p. 2025.03.22.644773). bioRxiv. 10.1101/2025.03.22.644773

20. Infanti, E., & Schwarzkopf, D. S. (2020). Mapping sequences can bias population receptive field estimates. NeuroImage, 211, 116636. 10.1016/j.neuroimage.2020.116636

21. Kok, P., Jehee, J. F. M., & de Lange, F. P. (2012). Less Is More: Expectation Sharpens Representations in the Primary Visual Cortex. Neuron, 75(2), 265–270. 10.1016/j.neuron.2012.04.034

22. Kok, P., Mostert, P., & de Lange, F. P. (2017). Prior expectations induce prestimulus sensory templates. Proceedings of the National Academy of Sciences, 114(39), 10473–10478. 10.1073/pnas.1705652114

23. Lee, K., & Rideaux, R. (2025). The influence of temporal context on vision over multiple time scales. eLife, 14. 10.7554/eLife.106614.1

24. Mostert, P., Kok, P., & de Lange, F. P. (2015). Dissociating sensory from decision processes in human perceptual decision making. Scientific Reports, 5(1), Article 1. 10.1038/srep18253

25. Oostenveld, R., & Praamstra, P. (2001). The five percent electrode system for high-resolution EEG and ERP measurements. Clinical Neurophysiology: Official Journal of the International Federation of Clinical Neurophysiology, 112(4), 713–719. 10.1016/s1388-2457(00)00527-7

26. Pelli, D. G. (1997). The VideoToolbox software for visual psychophysics: Transforming numbers into movies. 10.1163/156856897X00366

27. Press, C., Kok, P., & Yon, D. (2020). The Perceptual Prediction Paradox. Trends in Cognitive Sciences, 24(1), 13–24. 10.1016/j.tics.2019.11.003

28. Rao, R. P. N., & Ballard, D. H. (1999). Predictive coding in the visual cortex: A functional interpretation of some extra-classical receptive-field effects. Nature Neuroscience, 2(1), Article 1. 10.1038/4580

29. Richter, D., Heilbron, M., & de Lange, F. P. (2022). Dampened sensory representations for expected input across the ventral visual stream. Oxford Open Neuroscience, 1, kvac013. 10.1093/oons/kvac013

30. Rideaux, R. (2024). Task-related modulation of event-related potentials does not reflect changes to sensory representations. Imaging Neuroscience, 2, 1–13. 10.1162/imag_a_00266

31. Rideaux, R., Bays, P. M., & Harrison, W. J. (2025). Reply to: “Model mimicry limits conclusions about neural tuning and can mistakenly imply unlikely priors.” Nature Communications, 16(1), 6229. 10.1038/s41467-025-60860-9

32. Rideaux, R., Dang, P., Jackel-David, L., Buhmann, Z., Rangelov, D., & Mattingley, J. B. (2025). Violated predictions enhance the representational fidelity of visual features in perception. Journal of Vision, 25(4), 14. 10.1167/jov.25.4.14

33. Rideaux, R., West, R. K., Rangelov, D., & Mattingley, J. B. (2023). Distinct early and late neural mechanisms regulate feature-specific sensory adaptation in the human visual system. Proceedings of the National Academy of Sciences, 120(6), e2216192120. (world). 10.1073/pnas.2216192120

34. Rohenkohl, G., Gould, I. C., Pessoa, J., & Nobre, A. C. (2014). Combining spatial and temporal expectations to improve visual perception. Journal of Vision, 14(4), 8. 10.1167/14.4.8

35. Sasaki, Y., Rajimehr, R., Kim, B. W., Ekstrom, L. B., Vanduffel, W., & Tootell, R. B. H. (2006). The Radial Bias: A Different Slant on Visual Orientation Sensitivity in Human and Nonhuman Primates. Neuron, 51(5), 661–670. 10.1016/j.neuron.2006.07.021

36. Sereno, M., Dale, A., Reppas, J., Kwong, K., Belliveau, J., Brady, T., Rosen, B., & Tootell, R. (1995). Borders of multiple visual areas in humans revealed by functional magnetic resonance imaging. Science, 268(5212), 889–893. 10.1126/science.7754376

37. Sereno, M. I., Dale, A. M., Reppas, J. B., Kwong, K. K., Belliveau, J. W., Brady, T. J., Rosen, B. R., & Tootell, R. B. H. (1995). Borders of multiple visual areas in humans revealed by functional magnetic resonance imaging. Science, 268(5212), 889–893. 10.1126/science.7754376

38. Soltani, A., & Izquierdo, A. (2019). Adaptive learning under expected and unexpected uncertainty. Nature Reviews Neuroscience, 20(10), Article 10. 10.1038/s41583-019-0180-y

39. Stehr, D. A., Garcia, J. O., Pyles, J. A., & Grossman, E. D. (2023). Optimizing multivariate pattern classification in rapid event-related designs. Journal of Neuroscience Methods, 387, 109808. 10.1016/j.jneumeth.2023.109808

40. Summerfield, C., Trittschuh, E. H., Monti, J. M., Mesulam, M.-M., & Egner, T. (2008). Neural repetition suppression reflects fulfilled perceptual expectations. Nature Neuroscience, 11(9), 1004–1006. 10.1038/nn.2163

41. Tang, M. F., Kheradpezhouh, E., Lee, C. C. Y., Dickinson, J. E., Mattingley, J. B., & Arabzadeh, E. (2023). Expectation violations enhance neuronal encoding of sensory information in mouse primary visual cortex. Nature Communications, 14(1), Article 1. 10.1038/s41467-023-36608-8

42. Tang, M. F., Smout, C. A., Arabzadeh, E., & Mattingley, J. B. (2018). Prediction error and repetition suppression have distinct effects on neural representations of visual information. eLife, 7, e33123. 10.7554/eLife.33123

43. Teufel, C., Dakin, S. C., & Fletcher, P. C. (2018). Prior object-knowledge sharpens properties of early visual feature-detectors. Scientific Reports, 8(1), 10853. 10.1038/s41598-018-28845-5

44. Teufel, C., & Fletcher, P. C. (2020). Forms of prediction in the nervous system. Nature Reviews Neuroscience, 21(4), 231–242. 10.1038/s41583-020-0275-5

45. Whitaker, D., McGraw, P. V., & Levi, D. M. (1997). The influence of adaptation on perceived visual location. Vision Research, 37(16), 2207–2216. 10.1016/S0042-6989(97)00030-8

46. Wolpert, D. M., & Flanagan, J. R. (2001). Motor prediction. Current Biology, 11(18), R729–R732. 10.1016/S0960-9822(01)00432-8

47. Yon, D., Gilbert, S. J., de Lange, F. P., & Press, C. (2018). Action sharpens sensory representations of expected outcomes. Nature Communications, 9(1), 4288. 10.1038/s41467-018-06752-7

48. Zar, J. H. (1999). Biostatistical analysis. Pearson Education India.

